# Analysis of Spounaviruses as a Case Study for the Overdue Reclassification of Tailed Bacteriophages

**DOI:** 10.1101/220434

**Authors:** Jakub Barylski, François Enault, Bas E. Dutilh, Margo B.P. Schuller, Robert A. Edwards, Annika Gillis, Jochen Klumpp, Petar Knezevic, Mart Krupovic, Jens H. Kuhn, Rob Lavigne, Hanna M. Oksanen, Matthew B. Sullivan, Johannes Wittmann, Igor Tolstoy, J. Rodney Brister, Andrew M. Kropinski, Evelien M. Adriaenssens

**Author notes:** This paper is dedicated to Hans-Wolfgang Ackermann, a pioneer of prokaryotic virus electron microscopy and taxonomy, who died on February 12^th^, 2017, at the age of 80. He was involved in the early stages of this study, and his input is dearly missed.

## Abstract

It is almost a cliché that tailed bacteriophages of the order *Caudovirales* are the most abundant and diverse viruses in the world. Yet, their taxonomy still consists of a single order with just three families: *Myoviridae*, *Siphoviridae*, and *Podoviridae*. Thousands of newly discovered phage genomes have recently challenged this morphology-based classification, revealing that tailed bacteriophages are genomically even more diverse than once thought. Here, we evaluate a range of methods for bacteriophage taxonomy by using a particularly challenging group as an example, the Bacillus phage SPO1-related viruses of the myovirid subfamily *Spounavirinae*. Exhaustive phylogenetic and phylogenomic analyses indicate that the spounavirins are consistent with the taxonomic rank of family and should be divided into at least five subfamilies. This work is a case study for virus genomic taxonomy and the first step in an impending massive reorganization of the tailed bacteriophage taxonomy.

---

By the end of 2017, 3,033 complete genomes of tailed phages were available in the National Center for Biotechnology Information (NCBI) RefSeq database and a further 18,753 partial genomes were found in International Nucleotide Sequence Database Collaboration databases (Karsch-Mizrachi et al. 2012; O’Leary et al. 2016). The classification of this massive group is the formal responsibility of the Bacterial and Archaeal Viruses Subcommittee of the International Committee on the Taxonomy of Viruses (ICTV). In recent years, we (the Subcommittee) focused on classifying newly described phages into species and genera, within established viral families (Lavigne et al. 2008, 2009, Adriaenssens et al. 2015, 2017; Krupovic et al. 2016). However, once our attention shifted towards higher order relationships, we found that the ranks available in virus taxonomy (*species*, *genus*, *subfamily*, *family*, and *order*) were no longer sufficient for the description of phage diversity. The limitation is particularly acute in the case of the order *Caudovirales*—the most abundant and diverse group of viruses (Paez-Espino et al. 2016; Roux et al. 2016; Nishimura et al. 2017). Indeed, the diversity of caudovirads surpasses that of any other virus taxon. A recent analysis of the dsDNA virosphere using a bipartite network approach, whereby viral genomes are connected via shared gene families, demonstrated that the global network of dsDNA viruses consists of at least 19 modules, 11 of which correspond to caudovirads (Iranzo et al. 2016). Each of eight remaining modules encompasses one or more families of eukaryotic or archaeal viruses. Consequently, each of the 11 caudovirad modules could be considered a separate family. Despite this remarkable diversity, all caudovirads are currently classified into three families - *Myoviridae, Podoviridae*, and *Siphoviridae*. These families were historically established on morphological features alone, forming an artificial classification ceiling.

In this study, the Subcommittee explored the diversity of the order *Caudovirales* on the example of the *Spounavirinae* subfamily, a large group of myoviruses that forms one of the above-mentioned caudovirad modules (Iranzo et al. 2016; Bolduc et al. 2017). The subfamily was proposed in 2009 by Lavigne et al. to harbor Bacillus phage SPO1, Staphylococcus phage Twort, Staphylococcus phage K, Staphylococcus phage G1, Listeria phage P100, and Listeria phage A511 (Lavigne et al. 2009). The unifying characteristics of members of this subfamily are: the host belongs to the bacterial phylum *Firmicutes;* strictly virulent lifestyle; myovirion morphology; terminally redundant, non-permuted dsDNA genome 127–157 kb in length; and “considerable amino acid homology” (Klumpp et al. 2010). The strictly virulent lifestyle of these viruses has been somewhat disputed (Schuch and Fischetti 2009; Yuan et al. 2015) but still remains a rule of thumb for the taxon inclusion. Since the inception of the subfamily, the number of its members has grown significantly, and its taxonomic structure was contested several times (Klumpp et al. 2010; Barylski et al. 2014; Iranzo et al. 2016; Krupovic et al. 2016; Adriaenssens et al. 2017; Bolduc et al. 2017). At present, the *Spounavirinae* subfamily includes six genera (*Kayvirus, P100virus, Silviavirus, Spo1virus, Tsarbombavirus and Twortvirus*) and three unassigned species (*Enterococcus virus phiEC24C, Lactobacillus virus Lb338-1 and Lactobacillus virus LP65*).

Here, we reevaluated the current classification of spounavirins and related viruses and outlined a better fitting scheme, in the process also reaffirming the need for major changes in phage taxonomy that will better accommodate the observed genomic diversity.

## MATERIALS & METHODS

### Creation of the Dataset

Genome sequences of known spounavirins and spouna-like viruses were retrieved from GenBank or (preferably) RefSeq databases based on literature data, ICTV and taxonomic classifications provided by the NCBI. Records representing genomes of candidate spounavirins were retrieved by searching the same databases with the tBLASTx algorithm using as a queries terminase and major capsid proteins of Bacillus phage SPO1, Staphylococcus phage Twort, Bacillus phage Bastille, Listeria phage A511, Enterococcus phage φEF24C, and Lactobacillus phage LP65 [type isolates of the original subfamily, (Altschul et al. 1990; Brister et al. 2015)]. Sequences were manually curated and preclustered using Cluster Analysis of Sequences (CLANS; E-value cut-off 1e-10) to confirm their spounaviral affiliation (Frickey and Lupas 2004). This search yielded a set of 93 complete virus genomes, which were used in the following analyses (Supplementary Table 1).

The genomes were re-annotated using PROKKA with the settings --kingdom Viruses, --E-value 1e-6 (Seemann 2014). All original genome sequences are available from NCBI (accession number information listed in Supplementary Table 1) and the reannotated genomes from Github (github.com/evelienadri/herelleviridae).

### Genome-based Analyses

Gegenees (Ågren et al. 2012) was used to analyze genome similarities (fragment length 200 bp; step length 100 bp). Pairwise identities between all genomes under study were determined using BLASTn and tBLASTx algorithms with default parameters (Camacho et al. 2009). Symmetrical identity scores (% SI) were calculated for each pairwise comparison using the formula:

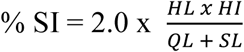

in which the HL is defined as the hit length of the BLAST hit, HI is defined as the percentage hit identity, QL is defined as the query length, and SL is defined as the subject length.

Symmetrical identity scores were converted into distances using the formula:

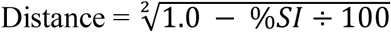

The resulting distance matrix was hierarchically clustered (complete linkage) using the hclust function of R (Development Core Team 2008). Trees were visualized using Itol (Letunic and Bork 2007).

Additionally, pairwise comparisons of the nucleotide sequences using VICTOR, a Genome-BLAST Distance Phylogeny (GBDP) method, were conducted under settings recommended for prokaryotic viruses (Meier-Kolthoff et al. 2014; Meier-Kolthoff and Göker 2017). The resulting intergenomic distances (including 100 replicates each) were used to infer a balanced minimum evolution tree with branch support via FASTME including subtree pruning and regrafting post-processing (Lefort et al. 2015) for each of the formulas D0, D4, and D6, respectively. Trees were visualized with FigTree (Rambaud 2007). Taxon demarcations at the species, genus and family rank were estimated with the OPTSIL program (Göker et al. 2009), the recommended clustering thresholds (Meier-Kolthoff and Göker 2017), and an F value (fraction of links required for cluster fusion) of 0.5 (Meier-Kolthoff et al. 2014).

### Proteome-based Analyses

The Phage Proteomic Tree was constructed as described previously (Rohwer and Edwards 2002) and detailed at https://github.com/linsalrob/PhageProteomicTree/tree/master/spounavirus. Briefly, the protein sequences were extracted and clustered using BLASTp. These clusters were refined by Smith-Waterman alignment using CLUSTALW version 2 (Larkin et al. 2007). Alignments were scored using open-source PROTDIST from the phylogeny inference package (PHYLIP) (Felsenstein 1989). Alignment scores were averaged and weighted as described previously (Rohwer and Edwards 2002) resulting in the final tree.

Orthologous protein clusters (OPCs) were constructed using GET_HOMOLOGUES software, which utilizes several independent clustering methods (Contreras-Moreira and Vinuesa 2013). To capture as many evolutionary relationships as possible, a greedy COGtriangles algorithm was applied with a 50% sequence identity threshold, 50% coverage threshold, and an E-value cut-off equal to 1e-10 (Kristensen et al. 2010). The results were converted into an orthologue matrix with the “compare_clusters” script (part of the GET_HOMOLOGUES suite) (Felsenstein 1989).

The OPCs defined above were used to compute the genomic fluidity for each pair of genomes. For two genomes i and j:

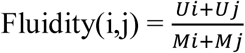

with Ui being the number of genes of i not found in j and Mi being the number of genes in i (Kislyuk et al. 2011). The resulting distance matrix was hierarchically clustered (complete linkage) using the hclust function of R (Development Core Team 2008). Trees were visualized using Itol (Letunic and Bork 2007).

Multiple alignments were generated for each OPC using Clustal Omega (Sievers et al. 2011). For each cluster, the amino acid identity between all protein pairs inside a cluster was determined using multiple alignment. For all genome pairs, the AAI (Konstantinidis and Tiedje 2005) was then computed and transformed into distance using the formula:

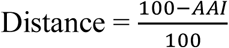

The resulting distance matrix was clustered and visualized as described above.

OPCs and multiple alignments for each cluster were used to determine a distance similar to the distance used to generate the Phage Proteomic Tree. To estimate protein distances, in this case, the distance (dist) function of the seqinR package (Charif et al. 2005)was preferred to PROTDIST of the PHYLIP package (Felsenstein 1989) as the resulting distances are between 0 and 1. Proteomic distances were then computed using the same formula as for the Phage Proteomic Tree. The results were clustered and visualized as described above.

The Dice score is based on reciprocal BLAST searches between all pairs of genomes A and B (Mizuno et al. 2013). The total summed bit-scores of all tBLASTx hits with ≥30% identity, alignment length ≥30 amino acids, and E-value ≤0.01 was converted to a distance DAB as follows:

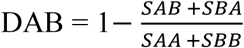

in which SAB and SBA represent the summed bit-scores between tBLASTx searches of A versus B, and B versusA, respectively, while SAA and SBB represent the summed tBLASTx bit-scores of the self-queries of A and B, respectively. The resulting distance matrix was clustered with BionJ (Gascuel 1997).

To investigate a genomic synteny-based classification signal, we developed a gene order-based metric built on dynamic programming, the Gene Order Alignment Tool (GOAT, Schuller et al.: Python scripts are available on request, manuscript in preparation). GOAT first identified protein-coding genes in the 93 spounavirin and spouna-like virus genomes using Prodigal V2.6.3 in anonymous mode (Hyatt et al. 2010), and assigned them to the latest pVOGs (Grazziotin et al. 2017)). pVOG alignments (9,518) were downloaded (http://dmk-brain.ecn.uiowa.edu/pVOGs/) and converted to profiles of hidden Markov models (HMM) using HMMbuild (HMMer 3.1b2, (Finn et al. 2011)). Proteins were assigned to pVOGs using HMMsearch (E-value <10-2) and used to generate a synteny profile of every genome. GOAT accounted for gene replacements and distant homology by using an all-vs-all similarity matrix between pVOG pairs based on HMM-HMM similarity (HH-suite 2.0.16) (Söding et al. 2005)). Distant HHsearch similarity scores between protein families were calculated as the average of reciprocal hits and used as substitution scores in the gene order alignment. The GOAT algorithm identified the optimal gene order alignment score between two virus genomes by implementing semi-global dynamic programming alignment based only on the order of pVOGs identified on every virus genome. To account for virus genomes being cut at arbitrary positions during sequence assembly, GOAT transmutes the gene order at all possible positions and in both sense and antisense directions in search of the optimal alignment score. The optimal GOAT alignment score GAB between every pair of virus genomes A and B, was converted to a distance DAB as follows:

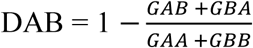

in which GAB and GBA represent the optimal GOAT score between A and B, and B and A, respectively, while GAA and GBB represent the GOAT scores of the self-alignments of A and B, respectively. This pairwise distance matrix was clustered with BionJ (Gascuel 1997).

Prokka re-annotated genomes were used to create pan-, core-, and accessory genomes of all selected spounavirins and spouna-like viruses (Seemann 2014). The annotations were analyzed using Roary (Page et al. 2015) with a 50% length BLASTp identity threshold for homologous genes. Roary functions as follows: CD-HIT (Fu et al. 2012) was used to precluster protein sequences and perform an all-vs-all comparison of protein sequences with BLASTp to identify orthologs and paralogs within the genomes. Markov cluster algorithm (MCL) (Enright et al. 2002) was then used to cluster the genomes based on the presence and absence of the accessory genes. The gene presence-absence output table from Roary was then imported into R and pairwise shared gene contents were calculated for each combination of genomes using a custom R-script (available from github.com/evelienadri/herelleviridae/tree/master). The resulting tree file was visualized using FigTree v1.4.3 (Rambaud 2007).

### Single Protein Phylogenies

Based on the OPC and pVOG analyses, we chose nine well-annotated protein clusters present in all 93 spounavirins and spouna-like viruses. Selected clusters included: DNA helicases, major capsid proteins, tail sheath proteins, two different groups of baseplate proteins, and four clusters with no known function. The members of these clusters were aligned using Clustal Omega with default parameters (Sievers et al. 2011). Resulting alignments were analyzed with ProtTest 3.4 (Darriba et al. 2011) to determine a suitable protein evolution model (only variations of models compatible with downstream software like JTT (Jones et al. 1992) and WAG (Whelan and Goldman 2001) were considered). Estimated models were used to generate phylograms with FastTree 2.1.7 (Price et al. 2010). The program implements the approximately maximum-likelihood method with Shimodaira-Hasegawa tests to generate the tree and calculate support of the splits. This approach is much faster than “traditional” maximum-likelihood methods with negligible accuracy loss (Price et al. 2010; Darriba et al. 2011; Liu et al. 2011).

## RESULTS

### General Overview

To determine the phylogenetic relationship between 93 known and alleged spounavirins, we used genomic, proteomic and marker gene-based comparative strategies. Regardless of the adopted phylogenetic approach applied, five separate, clear-cut clusters were identified. We believe that these clusters have a common origin and ought to come together under one umbrella taxon. We suggest to name this taxon “*Herelleviridae*,” in honor of the 100th anniversary of the discovery of prokaryotic viruses by Félix d’Hérelle (Table 1, Figs. 1-3 and Supplementary Table 1). The first cluster (here suggested to retain the name *Spounavirinae*) groups *Bacillus*-infecting viruses that are similar to Bacillus phage SPO1. The second cluster includes Bacillus-infecting viruses that resemble phage Bastille instead (named “*Bastillevirinae*” after the type species (Barylski et al. 2014)). The third cluster (“*Brockvirinae*,” named in honor of Thomas D. Brock, a microbiologist known for discovery of hyperthermophiles who worked on *Streptococcus* phages early in his career) comprises currently unclassified viruses of enterococci that are similar to Enterococcus phage φEF24C. The fourth cluster (“*Twortvirinae*,” named in honor of Frederick William Twort, the bacteriologist who discovered prokaryotic viruses in 1915) gathers staphylococci-infected viruses that are similar to Staphylococcus phage Twort. The remaining cluster (“*Jasinskavirinae*,” named in honor of Stanisława Jasińska-Lewandowska who was one of the first to study *Listeria* and its viruses) consists of viruses infecting *Listeria* that are similar to Listeria phage P100. The classification in five clusters left three viruses unassigned at this rank: Lactobacillus phage Lb338, Lactobacillus phage LP65, and Brochothrix phage A9.

These robust clusters can be further subdivided into smaller clades that correspond well with the currently accepted genera. The evidence supporting this suggested taxonomic reclassification is presented in the following sections.

### Genome-based Analyses

BLASTn analysis revealed that the genomes of several viruses were similar enough to consider them strains of the same species (they shared >95% nucleotide identity, Supplementary Fig. 1). The Staphylococcus viruses fell into four distinct, yet closely related groups corresponding to the established genera Twortvirus, Sep1virus, Silviavirus, and Kayvirus (Supplementary Fig. 1). With the exception of Enterococcus phage EFDG1, all Enterococcus viruses clustered as a clade representing a new genus (here suggested to be named “Kochikohdavirus” after the place of origin of the type virus of the clade, Enterococcus phage φEF24C (Uchiyama et al. 2008a, 2008b)). The Bacillus viruses clustered into the established genera Spo1virus, Cp51virus, Bastillevirus, Agatevirus, B4virus, Bc431virus, Nit1virus, Tsarbombavirus, and Wphvirus, with three species remaining unassigned at the genus rank (Table 1). These results were also confirmed with the Virus Classification and Tree Building Online Resource (VICTOR), a genome-BLAST distance phylogeny (GBDP) method (Supplementary Fig. 2) (Meier-Kolthoff and Göker 2017) and the Dice score (Supplementary Fig. 3), a tBLASTx-based measure that compares whole genome sequences at the amino acid level (Mizuno et al. 2013).

The patterns coalesced at a higher taxonomic level when the genomes were analyzed using tBLASTx (Supplementary Fig. 4). The *Enterococcus* viruses clustered into a single group sharing 41% genome identity, whereas the *Bacillus* viruses fell into two major groups, a group combining the genera *Spo1virus* and *Cp51virus*, and the remainder. All *Staphylococcus* viruses clustered above ≈36% genome identity, whereas *Listeria* viruses grouped with more than 79% genome identity. Overall, all these genomes were related at the level of at least 15% genome identity. *Lactobacillus* and *Brochothrix* viruses remained genomic orphans, peripherally related to the remainder of the viruses in this assemblage.

**Table 1.**
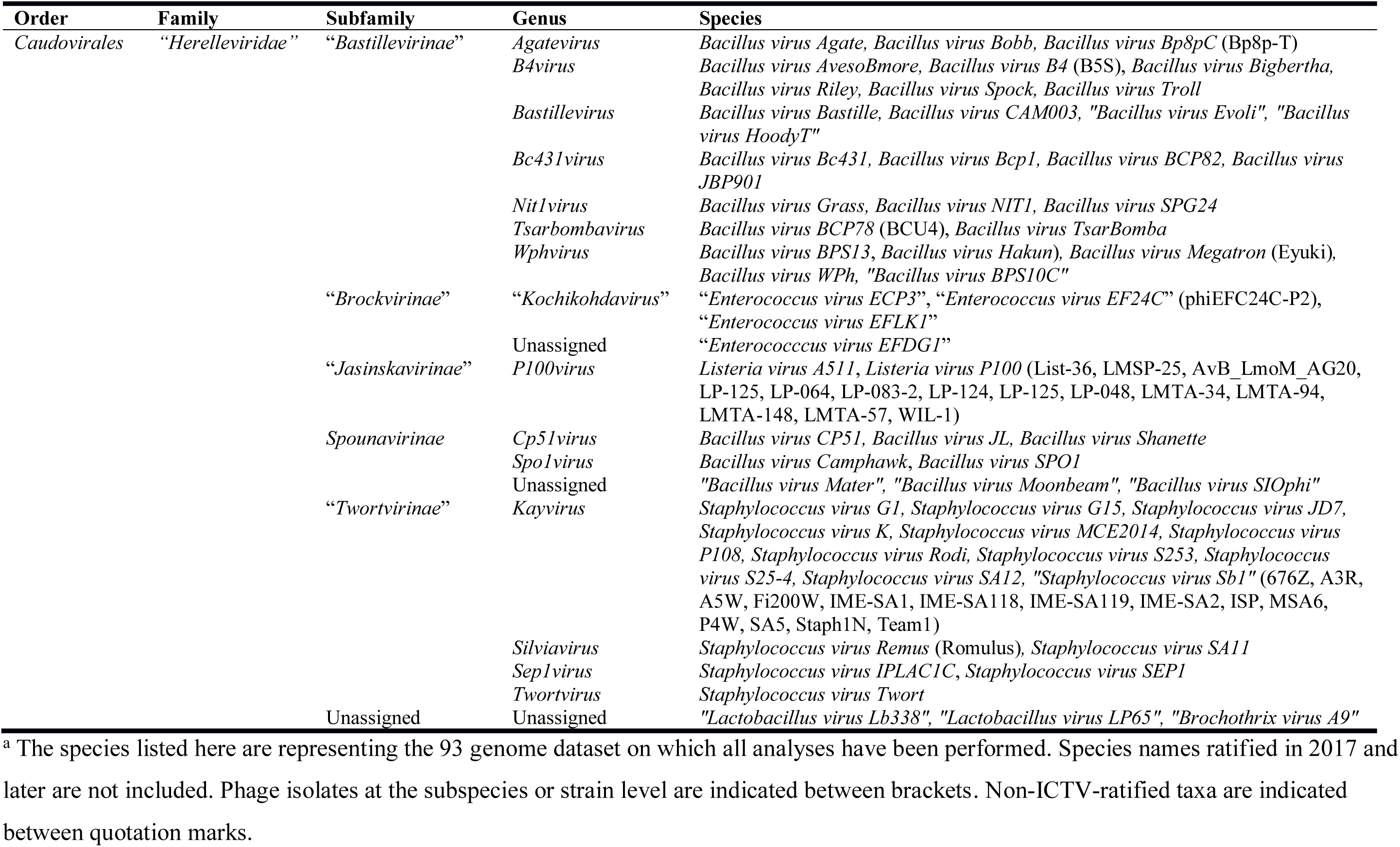
Suggested new classification of the 93 spounavirins and spouna-like viruses in the new family “Herelleviridae.”^a^

**Figure 1.**
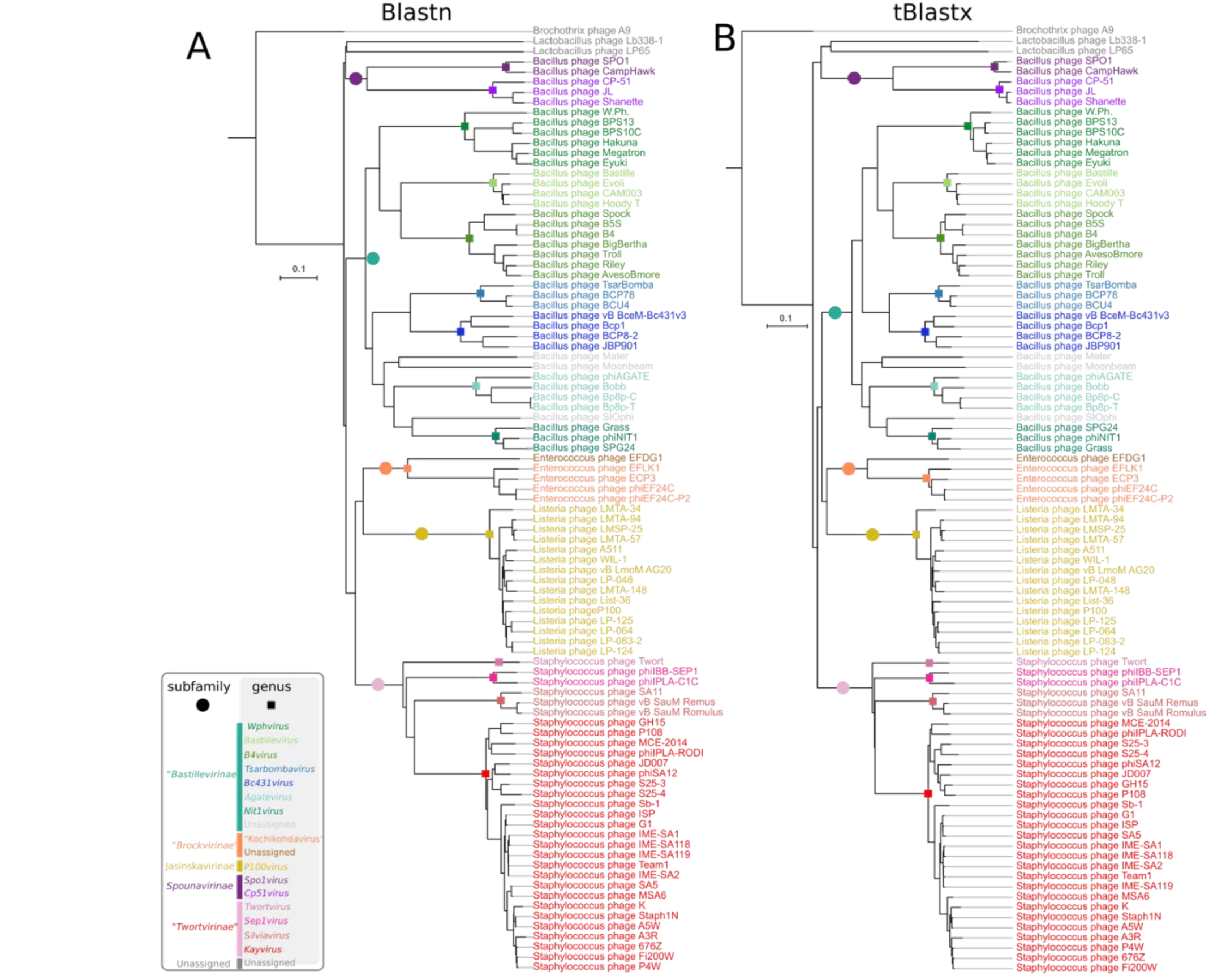
Genome-based clustering trees of 93 spounavirin and spouna-like viruses. A) Clustering was performed using nucleotide similarities (BLASTn) or B) translated nucleotide similarities (tBLASTx,). Genomes were compared in a pairwise fashion using Gegenees, transformed into a distance matrix, clustered using R, and visualized as trees using Interactive tree of life (Itol). The trees were rooted at Brochotrix phage A9. Genera and suggested subfamilies are delineated with colored squares and colored circles, respectively.

### Proteome-based Analyses

The virus proteomic tree showed five robust groupings corresponding with the suggested subfamilies (Fig. 2). Viruses that infect *Bacillus* fell into two groups as described before, represented by the revised *Spounavirinae* subfamily and the suggested new subfamily “*Bastillevirinae*.” Similarly, the *Listeria* and *Staphylococcus* viruses formed their own clusters, “*Jasinskavirinae*” and “*Twortvirinae*”, respectively. This clustering suggests that the major *Bacillus, Listeria*, and *Staphylococcus* virus groups are represented, but that further representatives are required from the under-sampled groups. The suggested “*Brockvirinae*” subfamily is under-sampled, and the grouping observed in the tree was not as well-supported as the other clusters.

Among 1,296 singleton proteins and 2,070 protein clusters defined using the orthologous protein clusters (OPC) approach, we identified 12 clusters common for all viruses (Table 2, Supplementary Table 2). Classification of the viral proteins using prokaryotic virus orthologous groups (pVOGs) showed that 38 pVOGs were shared between all 93 virus genomes (Table 2, Supplementary Table 3). This finding was in stark contrast with the results from core genome analysis using Roary, which revealed only one core gene (the tail tube protein gene). Upon closer inspection of the gene annotations, we found that these analyses might have been confounded by the presence of introns and inteins in many of the core genes (Supplementary Figs. 5–6). Indeed, many genes of spounavirins and related viruses are invaded by mobile introns or inteins (Goodrich-Blair et al. 1990; Lavigne and Vandersteegen 2013). These gaps in coding sequences challenge gene prediction tools and introduce additional bias in similarity-based cluster algorithms.

**Figure 2.**
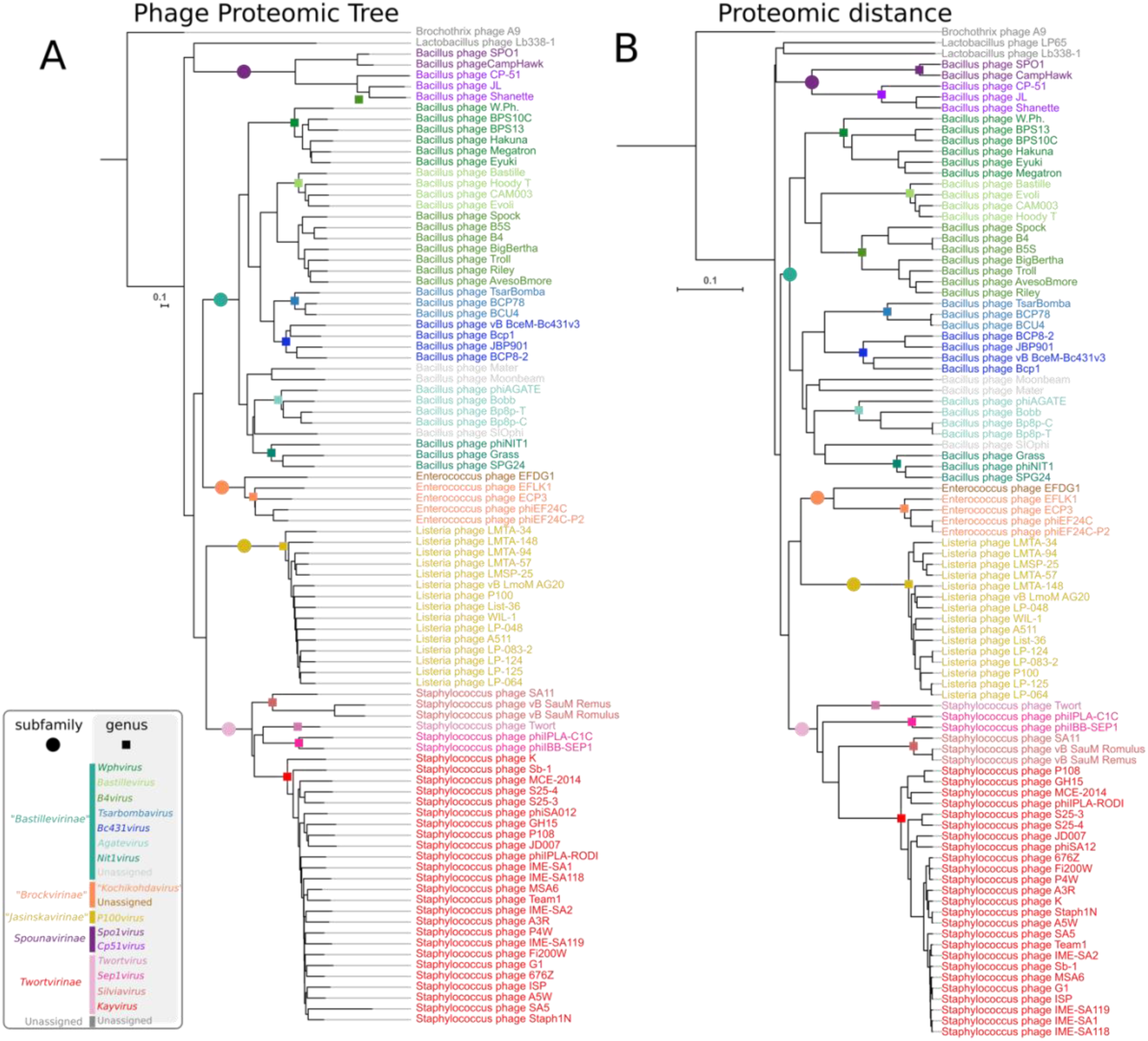
Predicted proteome-based clustering trees of 93 spounavirin and spouna-like viruses. A) Clustering was performed using the Phage Proteomics Tree approach and B) proteomic distance. Distances were calculated pairwise between all sets of predicted proteomes, clustered with R, and visualized using Itol. The trees were rooted at Brochotrix phage A9. Genera and suggested subfamilies are delineated with colored squares and colored circles, respectively.

The pairwise comparison of the predicted proteome content of the viruses revealed a very low overall similarity at the protein level (Supplementary Fig. 7). Most viruses shared less than 10% of their proteins. However, at the suggested new subfamily rank, we observed obvious virus groups sharing their proteomes. The *Enterococcus* viruses (“*Brockvirinae*”) shared over 35% of their protein content. The members of the *Bacillus* virus genera *Spo1virus* and *Cp51virus* of the subfamily *Spounavirinae* (*sensu stricto*) had approximately 20% of their proteins in common, whereas the *Bacillus* virus genera *Bastillevirus, B4virus, Bc431virus, Agatevirus, Nit1virus, Tsarbombavirus*, and *Wphvirus* (“*Bastillevirinae*”) and the *Staphylococcus* virus genera *Kayvirus, Silviavirus*, and *Twortvirus* (“*Twortvirinae*”) shared over 25% and over 30% of their predicted proteomes, respectively.

Genomic fluidity is a measure of the dissimilarity of genomes evaluated at the gene level (Kislyuk et al. 2011). Accordingly, the genomic fluidity results followed those obtained using proteome content analysis (Supplementary Fig. 8). Despite a high genomic fluidity for most of these viruses, the newly suggested subfamilies and genera were all supported.

The topology of the dendrogram obtained using the average amino acid identity (AAI) approach also supported the suggested new taxonomic scheme (Supplementary Fig. 8). The AAI was greater than 35% within each subfamily and greater than 67% within each genus. The AAI of all viruses analyzed in this study was not lower than 22%. The members of the genus *Wphvirus* had the lowest AAI (76%) and the lowest AAI for a pair of proteomes (67% between Bacillus phage W.Ph. and Bacillus phage Eyuki) but surprisingly they had a mid-range genomic fluidity (0.15), suggesting that the protein sequences of wphviruses might have evolved rapidly.

**Table 2.** Core genes with putative annotated functions identified in all 93 spounavirin and spouna-like virus genomes.

| Putative function of the core gene identified<sup>a</sup> | pVOG<sup>b</sup> / OPC<sup>b</sup> ID | Identification method |
| --- | --- | --- |
| DnaB-like helicase | VOG0025, OPC6121 | OPC, pVOG |
| Baseplate J-like protein | VOG4691, VOG4644, OPC6132 | OPC, pVOG |
| Tail sheath protein | VOG0067, OPC6142 | OPC, pVOG |
| Terminase large subunit | VOG0051, OPC6160 | pVOG |
| Major capsid protein | VOG0061, OPC6148 | OPC, pVOG |
| Prohead protease | VOG4568, OPC6150 | pVOG |
| Portal protein | VOG4556, OPC6151 | OPC, pVOG |
| DNA primase | VOG4551 | pVOG |
| DNA polymerase I | VOG0668, OPC6097 | OPC, pVOG |
| RNA polymerase | VOG0118 | pVOG |
| Recombination exonuclease | VOG4575 | pVOG |
| Recombination endonuclease | VOG0083 | pVOG |
| Tail tape measure protein | VOG0069 | pVOG |
| Tail tube protein | VOG0068, OPC6141 | OPC, pVOG, Roary |

<sup>a</sup> The full lists of orthologous proteins and pVOGs are available in Supplementary
Tables 2 and 3, respectively.
<sup>b</sup> pVOG, prokaryotic virus orthologous group; OPC, orthologous protein clusters.

The pangenome of the spounavirins and spouna-like viruses (4,182 genes) as calculated using Roary (Page et al. 2015) was further analyzed by clustering the genomes based on the presence or absence of the accessory genes (Supplementary Fig. 9). The obtained tree supported the current division of the viruses into approved genera and the suggested new subfamilies.

Many virus genomes are thought to be highly modular, with recombination and horizontal gene transfer potentially resulting in “mosaic genomes” (Juhala et al. 2000; Krupovic et al. 2011). By clustering the spounavirin and spouna-like virus genomes based solely on the gene order of their genomes, we investigated whether gene synteny was preserved (Supplementary Fig. 10). The results revealed that genomic rearrangements leave a measurable evolutionary signal in all lineages, since the genomic architecture analysis clustered all viruses according the suggested taxa. The potential exception was Bacillus phage Moonbeam (Cadungog et al. 2015). However, we did not observe the high modularity that may be expected with rampant mosaicism. The lack of considerable mosaicism supports the recent findings by Bolduc et al. that, at most, about 10% of reference virus genomes have a high degree of mosaicism (Bolduc et al. 2017). Thus, while the gene order in viruses belonging to the newly suggested family “*Herelleviridae*” is not necessarily strictly conserved, we observed a clear evolutionary pattern that is consistent with the sequence-based approaches tested in this study.

### Single Protein Phylogenies

The phylogenetic trees based on comparisons of the major capsid, tail sheath, and DnaB-like helicase proteins are presented in Figure 3. All nine phylograms based on OPC are included in the trees in Supplementary Figure 11. For nearly all single marker trees, the topologies supported the suggested taxonomic scheme. Generally, each taxon is represented as a separate branch on the dendrogram. Notable exceptions could be found in two trees based on hypothetical proteins (OPCs 10357 and 10386). The first protein places the revised subfamily *Spounavirinae* as a subclade of “*Bastillevirinae”* and the second protein shuffled viruses from the genera *Silviavirus* and *Kayvirus*. This result may indicate that some degree of horizontal gene transfer occurs between groups, which share common hosts.

**Figure 3.**
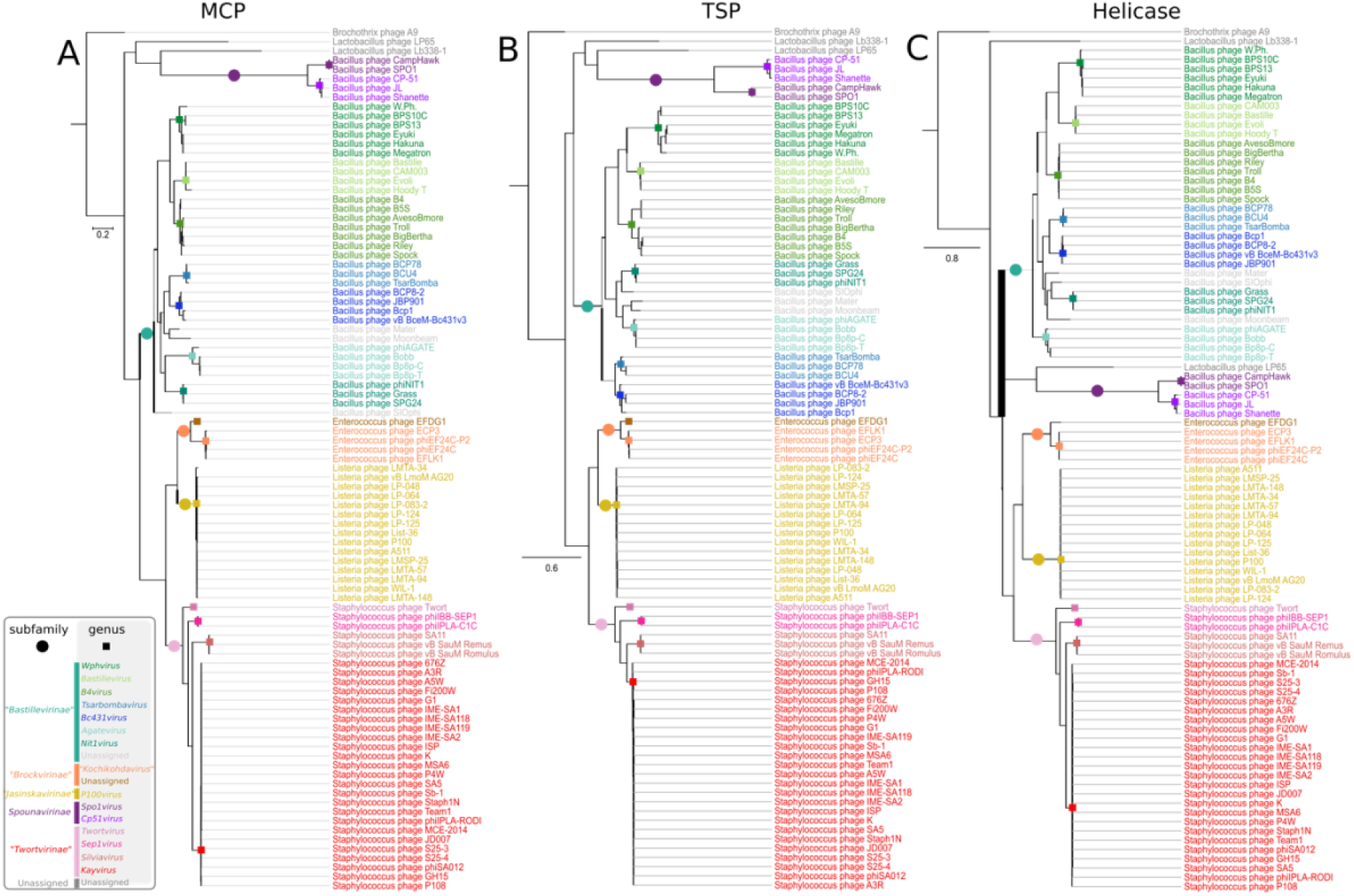
Phylogenetic trees based on comparisons of major protein clusters of amino acid sequences of spounavirin and spouna-like viruses. Amino acid sequences from A) the major capsid protein, B) tail sheath protein, and C) helicase were aligned with Clustal Omega, and trees were generated using FastTree maximum likelihood with Shimidaira-Hasegawa tests. The scale bar represents the number of substitutions per site. The trees were rooted at Brochotrix phage A9. Genera and suggested subfamilies are delineated with colored squares and colored circles, respectively.

## DISCUSSION

Taxonomic methods must constantly develop to keep up with the ever-increasing pace of virus discovery. In the rapidly expanding field of phage studies, this requirement proved to be problematic, and although there are more than 3000 publicly available caudovirad genomes, only 873 have been officially classified by the ICTV (Davison 2017). The remaining genomes are provisionally stashed within “unclassified” bins attached to the order or associated families (Adams et al. 2017; Simmonds et al. 2017). We believe that this work is an important step toward solving the problem of these “phage orphans”. This study represents the first example of a true taxonomic assessment from an “ensemble of methods”. We are encouraged that the combination of genome sequence analyses, virus proteomic trees, core protein clusters, gene order genomic synteny (GOAT), and single gene phylogenies yields consistent and complementary results. Convergence of the results reasserts the usefulness of genome-based classification at a higher taxonomic rank and the ability of these methods to accommodate viral diversity.

All evidence considered, we suggest that the spounavirins should be removed from the family *Myoviridae* and given a family rank. Hence, we propose establishing a new family “Herelleviridae”, in the order *Caudovirales* next to a smaller *Myoviridae* family. The new family would contain five subfamilies: *Spounavirinae* (sensu stricto), “*Bastillevirinae*”, “*Twortvirinae*”, “Jasinkavirinae”, and “*Brockvirinae*”, each comprising the genera listed in Table 1 (with additional information in S1 Table). The suggested classification corresponds well with the taxonomy of the hosts and leaves only 3% of viruses within the new family unassigned at the genus and subfamily rank. These unassigned viruses may represent clades at the genus and subfamily rank that are still under-sampled.

We believe that detachment of spounavirins from their original taxon will soon be followed by abolishment of the *Podoviridae, Myoviridae* and *Siphoviridae* families, in combination with the addition of new taxon ranks (e.g., class) required to accommodate the observed diversity of tailed phages. Substitution of the current families with a set of new “phylogenomic” ones will more faithfully reflect the genetic relationships of these viruses. This change does not remove the historically established virus morphotypes observed among caudovirads: myovirids forming particles with contractile tails, siphovirids forming particles with long non-contractile tails, and podovirids forming particles with short non-contractile ones. By disconnecting morphotype and family classification, taxonomically related clades can be grouped across different morphotypes. Such an approach would solve the problems of the muviruses that are suggested to be classified in the family “*Saltoviridae*” (Hulo et al. 2015) and potentially the broad set of Escherichia phage lambda-related viruses that are currently distributed among the families *Siphoviridae* and *Podoviridae* (Grose and Casjens 2014).

## ACKNOWLDEGMENTS

We thank Laura Bollinger, Integrated Research Facility at Fort Detrick, for technical writing services. E.M.A. would like to thank P.H. Nel for assistance with R scripting.

